# Neutron crystallography of the covalent intermediate of β-glucosidase reveals remodeling of the catalytic center

**DOI:** 10.1101/2025.02.06.636774

**Authors:** Naomine Yano, Hiromu Arakawa, Chih-Chieh Lin, Akihiro Ishiwata, Katsunori Tanaka, Katsuhiro Kusaka, Shinya Fushinobu

**Author notes:** Corresponding authors: Shinya Fushinobu, 1-1-1 Yayoi, Bunkyo-ku, Tokyo 113-8657, Japan; and Naomine Yano, 1-1-1, Kouto, Sayo-cho, Sayo-gun, Hyogo 679-5198 Japan., **Email:** and. **Author Contributions:** S.F. administrated the project; N.Y. and S.F. designed the research; N.Y., H.A., C.C.L., and K.K. performed the research; A.I. and K.T. provided the reagents; N.Y. and K.K. contributed analytics tools; N.Y., C.C.L., K.K., and S.F. analyzed the data; N.Y. and S.F. wrote the paper. **Competing Interest Statement:** The authors declare no competing interests.

## Abstract

Anomer-retaining glycoside hydrolases (GHs) generally catalyze a double displacement reaction via a covalent intermediate; however, neutron crystallography of glycoside ligand-bound states has not been performed. In this study, we investigated β-glucosidase Td2F2 from GH family 1 as a model enzyme for anomer-retaining GHs. We determined joint X-ray/neutron structures of Td2F2 in ligand-free form, covalent intermediate with a 2-deoxy-2-fluoro glucoside (2F-Glc) inhibitor, and glucose product complex using hydrogen/deuterium-exchanged crystals at room temperature, with neutron diffraction resolutions of 1.80–1.70 Å. Extensive hydrogen bonds recognizing the hydroxy groups of 2F-Glc were identified, along with the positions of deuterium atoms. The acid/base catalyst residue Glu166 was anchored by a hydrogen bond network pivoted by Asn293. Tyr295 forms a hydrogen bond with the catalytic nucleophile residue Glu352 in the ligand-free and glucose complex forms, while the active center undergoes significant reorganization, including side chain displacements of Glu352 and Tyr295, as well as the incorporation of a water molecule. An alternative conformation of Tyr295 was observed in the 2F-Glc structure at room temperature, suggesting its role in positioning the nucleophilic water during the deglycosylation step. The tyrosine hydrogen bonded to the nucleophile is also conserved in many other anomer-retaining GH families, underscoring its importance in catalysis. Based on the deuterium/hydrogen positions determined from neutron structures, we proposed a detailed reaction mechanism for Td2F2.

**Significance Statement:** Glycoside hydrolases perform diverse functions in organisms, with over 180 known enzyme families. Although hydrogen bonds and proton transfer play important roles in enzymatic reactions, hydrogen atoms are generally invisible in macromolecular X-ray crystallography. In anomer-retaining glycoside hydrolases, general acid/base catalysis and the formation and hydrolysis of a covalent glycosyl-enzyme intermediate have been postulated. Here, we report neutron crystal structures of a β-glucosidase, where hydrogen and deuterium atoms were visualized at high resolution. An intricate hydrogen-bonding network and remarkable remodeling at the catalytic center were observed during covalent intermediate formation, revealing a detailed catalytic mechanism. The enzyme belongs to glycoside hydrolase family 1 and represents numerous enzymes employing the anomer-retaining mechanism.

## Introduction

Glycoside hydrolases (GHs, EC 3.2.1.-) play essential roles in carbohydrate metabolism by significantly accelerating glycosidic bond cleavage, one of the most stable linkages in biological macromolecules (1, 2). GHs constitute the most prevalent enzyme class, with over 180 GH families currently listed in the Carbohydrate-Active enZymes (CAZy) database (3, 4). These enzymes employ two standard catalytic mechanisms involving two carboxylic acid (Asp or Glu) residues as catalytic groups: either retaining or inverting the α/β-anomeric configuration of sugar after hydrolysis (5). The anomer-retaining mechanism is more widespread than the inverting mechanism, utilized by over half of the GH families. Anomer-retaining β-glucosidase (BGL, EC 3.2.1.21) is a major exo-type GH that acts on a β-D-glucopyranoside bond at the non-reducing end of oligomeric (and sometimes polymeric) substrates, releasing β-glucose. Anomer-retaining BGLs belong to families GH1, GH2, GH3, GH4, GH5, GH30, GH39, and GH116 in CAZy (3). BGLs have been isolated from archaea, bacteria, fungi, plants, and environmental genomes, as they play key roles in natural carbohydrate utilization (6). BGL is also an important component of cellulase cocktails used in the industrial degradation of renewable plant biomass and has applications in the food industry (7, 8).

Unlike inverting-type GHs, which utilize a simpler single-displacement mechanism with general acid and base catalyst residues, retaining GHs, including BGLs, use a two-step double-displacement mechanism. In this mechanism, one residue acts as a nucleophile and the other as a general acid/base catalyst (see the Lexicon page in CAZypedia https://www.cazypedia.org/) (9). In the first glycosylation step (glycosidic bond cleavage and aglycone release), the nucleophile residue forms a covalent glycosyl-enzyme intermediate (GEI), while the acid/base catalyst residue donates a proton to the glycosidic bond oxygen. In the subsequent deglycosylation step (glycone release), the acid/base catalyst activates a nucleophilic water (NW) molecule by accepting a proton, facilitating GEI hydrolysis. Thus, proton transfer and hydrogen bonds are crucial for catalysis and substrate recognition in GHs, which specifically act on carbohydrates with numerous hydroxy groups. However, H atoms are generally not visible in widely used protein structure determination methods, such as X-ray crystallography and cryogenic electron microscopy. Neutron crystallography, using deuterium (D)-exchanged samples, is a promising technique for determining the positions of H and D atoms in proteins (10), but it has been conducted on a limited number of GHs. Despite more than 6400 GH structures (EC 3.2.1.-) being deposited in the Protein Data Bank (PDB), mainly determined by X-ray crystallography, neutron structures have been reported for only five enzymes: GH22 hen-egg white lysozyme (11–15), GH22 human lysozyme (16), GH24 T4 lysozyme (17, 18), GH11 xylanase (19), and GH45 endoglucanase (20). Additionally, only one ligand-bound neutron crystal structure has been reported for the GH45 endoglucanase from *Phanerochaete chrysosporium*, in which cellopentaose was bound at subsites from +1 to +5, and the catalytically relevant –1 subsite was unoccupied (20). A major challenge in neutron protein crystallography is the need for longer beam exposure times and larger crystal volumes (> 1 mm^3^) compared to those for X-ray crystallography, as both the beam intensity at available neutron source facilities and beam interaction with biological soft matter are significantly weaker (10).

In this study, we applied joint X-ray/neutron (XN) crystallography to a thermophilic, glucose-tolerant GH1 BGL (Td2F2) isolated from a compost metagenome (21). Our previous crystallographic study has shown that Td2F2 crystals grow large and diffract X-rays at high resolutions (∼ 1.1 Å) (22), making them suitable for neutron crystallography. In addition to the ligand-free (apo) and glucose complex structures, we determined a covalent intermediate structure using a 2-deoxy-2-fluoro glucoside (2F-Glc) inhibitor (23), which has been widely used for X-ray crystallography of GHs (24). The three high-resolution XN structures, containing D and H atoms at ambient temperature, revealed important structural features of Td2F2 related to the reactions and interactions at the –1 subsite of BGL during the catalytic cycle.

## Results and Discussion

### Diffraction experiments and joint XN structure determination

The recombinant Td2F2 protein produced in *Escherichia coli* was used for crystallization. H/D exchange was achieved by replacing the purified protein solution with a deuterated buffer, and crystals were grown using solutions prepared with heavy water containing CHES-NaOD buffer (pD 9.5). Glucose complex crystals were grown in the presence of 200 mM glucose. To obtain the covalent intermediate structure, the Td2F2 protein was incubated with 2,4-dinitrophenyl-2-deoxy-2-fluoro-β-D-glucopyranoside (DNP-2F-Glc) to form the 2F-Glc complex, which was then crystallized. Crystals grew to approximately 5.5, 3.0, and 4.8 mm^3^ in the ligand-free form, glucose complex form, and 2F-Glc complex form, respectively (*SI Appendix*, Figs. S1–S3). The crystals were used for multi-probe quantum beam diffraction experiments. The resolutions of the X-ray and neutron diffraction datasets collected at room temperature were 1.19 and 1.80 Å in the ligand-free form, 1.20 and 1.70 Å in the glucose complex form, and 1.32 and 1.70 Å in the 2F-Glc complex form, respectively (*SI Appendix*, Tables S1–S3). The X-ray data resolution collected in this study was improved compared to previously reported data for the ligand-free form (1.60 Å, PDB ID: 3WH5) and the glucose complex form (1.60 Å, PDB ID: 3WH6), which were collected at cryogenic temperatures (22). Non-hydrogen atoms were modeled according to the X-ray electron density (XRED) map. Joint XN structure refinements were successfully completed with reasonable statistics for crystallography and protein stereochemistry. In the final XN structure, 3,227 H atoms, 679 D atoms, and 346 water molecules in the ligand-free form, 3,278 H atoms, 815 D atoms, and 275 water molecules in the glucose complex form, and 3,269 H atoms, 865 D atoms, and 304 water molecules in the 2F-Glc complex form were modeled based on the neutron scattering length density (NSLD) maps. Alternative conformations of the main and side chains were not adopted because the H/D alternative conformer became even more alternative, and the occupancy could not be estimated correctly.

### Protonation states in the active site

Fig. 1 shows the NSLD maps in the active site of the ligand-free, glucose complex, and 2F-Glc complex forms. In the ligand-free structure, five water molecules were bound to subsite –1, and no positive D atom peaks were observed in any of them (Fig. 1*A*). Protonation of the hydroxy groups of glucose and 2F-Glc in the complex structures was determined based on the density peaks that appeared via H/D exchange. In the glucose complex, weak positive peaks were observed at the O2 and O3 hydroxy groups of glucose, with D atom occupancies of 0.54 to 0.52, respectively, while no H/D atom was observed at O1, O4, and O6 (Fig. 1*B* and *SI Appendix*, Fig. S4 and Table S4). In the 2F-Glc complex, strong positive peaks were observed at the O3, O4, and O6 hydroxy groups, with D atom occupancies ≥ 0.71 (Fig. 1*C* and *SI Appendix*, Table S4). The hydroxy groups of glucose and 2F-Glc form hydrogen bonds with surrounding protein residues, and strong positive peaks were clearly observed at the exchanged D atoms of these residues (Asn165, His121, Asn20, and Trp407), as well as at the aromatic stacking residue at subsite –1 (Trp399), especially in the 2F-Glc complex structure. Therefore, the recognition of the 2F-Glc intermediate by the enzyme could be determined in detail, including the H/D atoms.

**Fig. 1.**
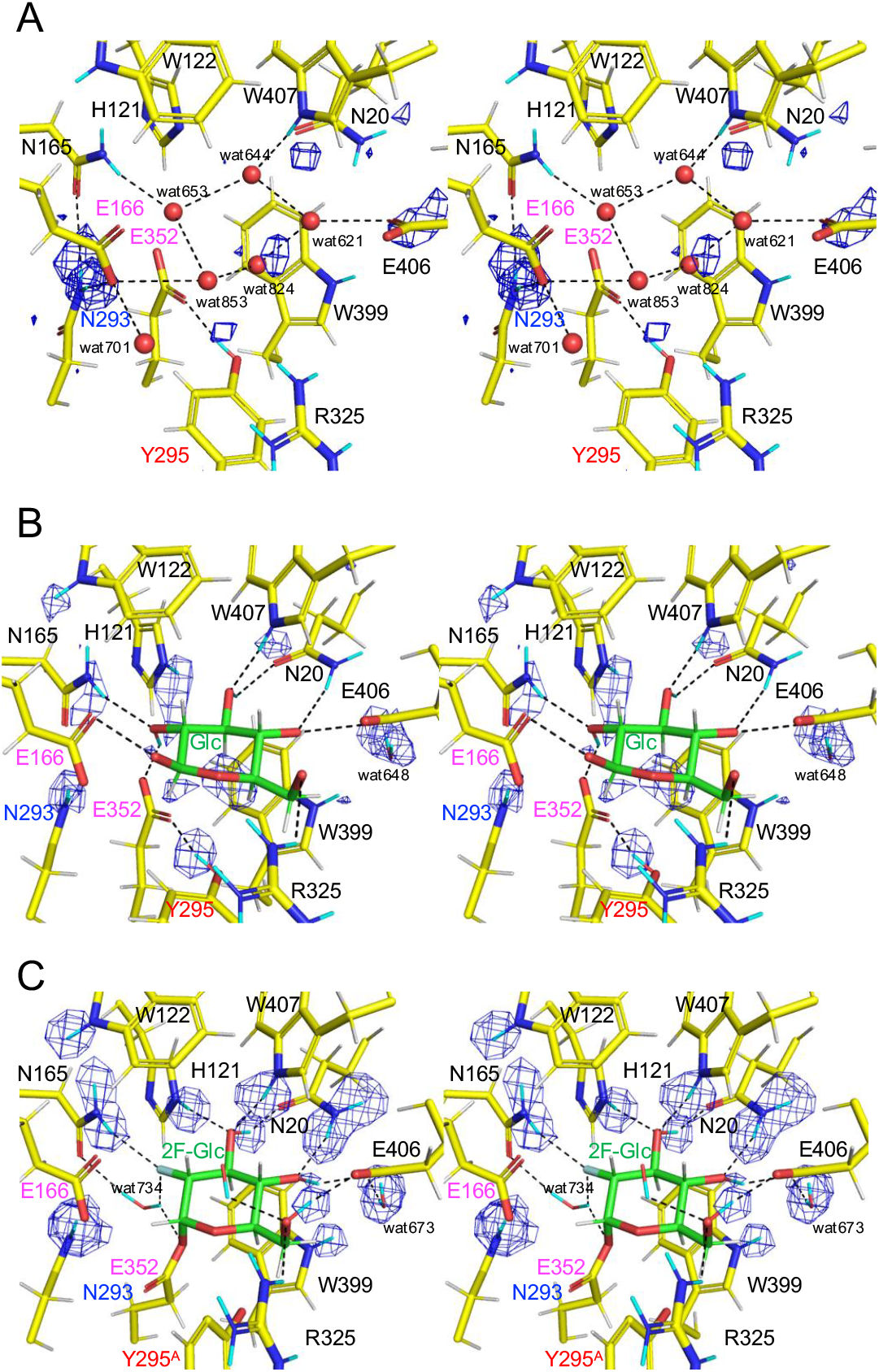
Stereo view of the active site with *mF*_o_-*DF*_c_ neutron scattering length density (NSLD) maps (blue mesh). (*A*) Ligand-free structure (2.9σ). (*B*) Glucose complex structure (3.0σ). (*C*) 2F-Glc complex structure (3.3σ). H and D atoms are colored white and cyan, respectively. Hydrogen bonds are shown with black dotted lines. The following atoms were excluded from the map calculation: DD21 and DD22 of Asn20, Asn165, and Asn293; DE1 of His121; DE1 of Trp122 and Trp399; DH1 of Tyr295; DE, DH11, DH12, DH21, and DH22 of Arg325. DO2 and DO3 of glucose and D1 and D2 of wat648 in (*B*); and DO3, DO4, and DO6 of 2F-Glc, and D1 and D2 of wat673 in (*C*) were also excluded from the map calculation.

No positive peak was observed near the carboxyl group of the acid/base catalyst residue Glu166 in any of the three states (Fig. 1). This was likely due to the high pD conditions (pD 9.5) for the growth of large Td2F2 crystals, which is higher than the optimal pH (pD = pH + 0.4). The activity of Td2F2 toward *p*-nitrophenyl-β-D-glucopyranoside (pNP-Glc) was optimal at pH 5.5 but reduced to approximately 5% at pH 8.52 (21). In contrast, two strong positive peaks were observed near the ND1 atom of Asn293 (DD21 and DD22), forming an evident hydrogen bond with Glu166 in all three states with D atom occupancy ≥ 0.60 (Fig. 2*A* and *SI Appendix*, Fig. S5 and Table S4). DD21, DD22, and OD2 atoms of Asn293 formed hydrogen bonds with the OD1 atom of Asn165, OE2 atom of Glu166, and DG1 atom of Thr351, respectively. Asn293 is buried inside the protein, with its ND2 atom surrounded by several residues (*SI Appendix*, Fig. S6). The extensive hydrogen bond network starting from the amide nitrogen of Asn293 anchors the side chains of Glu166 and Asn165 and extends to the active site, where several water molecules are fixed in the ligand-free form (Fig. 1*A*). Because Asn293 is not directly accessible from solvent water atoms, its D atoms were transferred from D_2_O via the OE2 atom of Glu166 through amide-imidic acid tautomerization of the Asn side chain. Therefore, the high H/D-exchanged state at the ND2 atom of Asn293 indirectly showed that the carboxyl group of Glu166 was transiently protonated even in an environment where the pD was above 9.0. The temperature factors of the O atom of wat701 and wat853 were ≥ 29 Å^2^, whereas those of wat653 and wat644 were ≤ 22 Å^2^ (*SI Appendix*, Table S4), indicating that wat701 and wat853 are more easily replaced by the substrate. The hydrogen bond network pivoted by Asn293 anchors Glu166 in the glucose and 2F-Glc complex structures as well (*SI Appendix*, Fig. S5). Therefore, this network is crucial for the function of the acid/base catalyst at a static position throughout the reaction cycle.

**Fig. 2.**
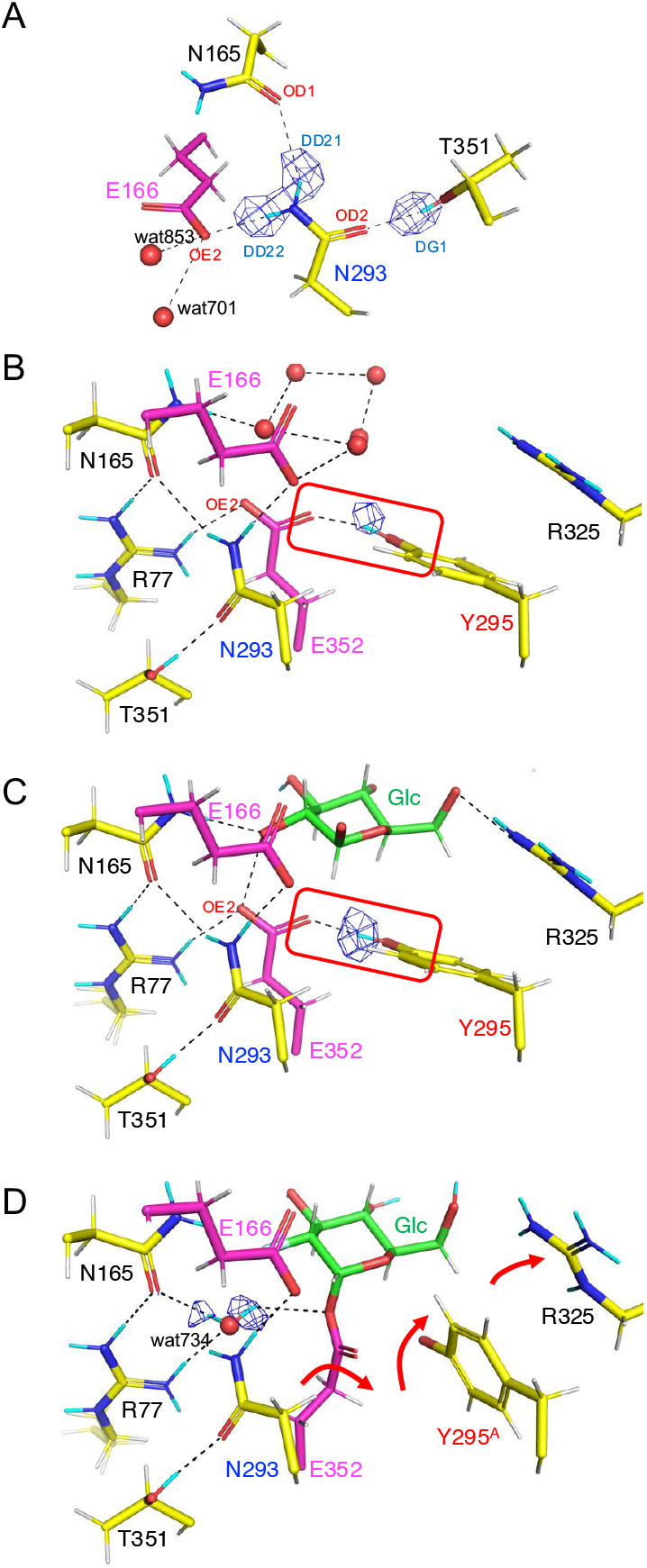
Structures around Asn293 and acid/base catalyst Glu166 with *mF*_o_-*DF*_c_ NSLD maps (blue mesh). (*A*) Asn293 and surrounding residues in the ligand-free structure (4.0σ). (*B*-*D*) The nucleophile Glu352, Tyr295, and surrounding residues in (*B*) ligand-free, (*C*) glucose complex, and (*D*) 2F-Glc complex structures (3.0σ). Red boxes in (B) and (C) indicate the hydrogen bond between Glu352 and Tyr295. Red arrows in (D) indicate the side chain displacements of Glu352, Tyr295, and Arg325. In the X-ray/neutron joint structure, a single conformation (Y295^A^) was placed in the coordinates. H and D atoms are colored white and cyan, respectively. Hydrogen bonds are shown with black dotted lines. The following atoms were excluded from the map calculation: DD21 and DD22 of Asn293 and DG1 of Thr351 in (*A*); DH of Tyr295 in (*B*) and (*C*); and D1 and D2 of wat734 in (*D*).

A clear NSLD peak at the side chain hydroxy of Tyr295 (D atom occupancy ≥ 0.53) was observed in the ligand-free and glucose complex structures (Fig. 2*BC* and *SI Appendix*, Table S4), and the side chains of Tyr295 and Glu352 form a hydrogen bond. Because Glu352 is the catalytic nucleophile residue, this hydrogen bond is crucial for the catalytic function of Td2F2. In most clan GH-A family enzymes (e.g., GH2, GH5, and GH17) (25), whose active centers are conserved with GH1, a tyrosine residue hydrogen bonded to the nucleophile glutamate is conserved (*SI Appendix*, Fig. S7). In the 2F-Glc complex structure, covalent bond formation induced side chain deviation of Glu352 from the other two structures (Fig. 2*D*), as previously observed in the X-ray crystal structures of other GH1 BGLs (26–28). The side chains of Tyr295 and Arg325 are also deviated, disrupting their hydrogen bonds with Glu352 and the O6 hydroxy group of glucose. Additionally, a new water molecule (wat734) was present in the 2F-Glc complex structure at a position corresponding to the OE2 atom of Glu352 in the ligand-free and glucose complex structures. Wat734 exhibited two positive NSLD peaks (D atom occupancy ≥ 0.39, *SI Appendix*, Table S4) and was stabilized by hydrogen bonds with Glu352, Asn165, and Arg77.

### Alternative conformation of a key residue for deglycosylation, Tyr295

In the XRED map of the 2F-Glc complex structure, an alternative conformation (B) in which Tyr295 moved significantly, including the main chain, was observed (Fig. 3*C*). In the XN joint structure, only conformer A was included in the coordinates for refinement, whereas alternative conformations were refined using X-ray diffraction data only (*SI Appendix*, Table S5). Tyr295 displayed a conformer ratio of A:B = 0.54:0.46. The side chain hydroxy of Tyr295^B^ was hydrogen-bonded to the acid/base catalyst Glu166 and was positioned 3.7 Å from the anomeric (C1) carbon of 2F-Glc. This alternative conformation was not observed in the ligand-free or glucose complex structures (Fig. 3*AB*). Interestingly, alternative conformation of tyrosine in the active center has never been observed in the crystal structures of GH1 enzymes (BGL and myrosinase) with diffraction data collected at cryogenic temperatures (*SI Appendix*, Table S6 and Fig. S8). When we flash-cooled a 2F-Glc complex crystal of Td2F2 and measured X-ray diffraction data at 100 K (*SI Appendix*, Table S7), no alternative conformation of Tyr295 was observed (Fig. 3*D*). These results indicate that the alternative conformation of tyrosine in the active center of GH1 BGL is only detectable in crystallographic diffraction data measured at room temperature. The position of the side chain hydroxy group of Tyr295^B^ suggests that this residue contributes to the deglycosylation step from the GEI. During deglycosylation, a NW attacks the anomeric carbon of the GEI, with deprotonation via the acid/base catalyst residue (5). Efficient S_N_2 nucleophilic attack requires approaching from the backside of the scissile bond. In anomer-inverting GHs, in addition to the base catalyst, another protein residue assists this nucleophilic attack by holding the NW by a hydrogen bond (29). In Fig. 3*E*, a putative NW was placed in the Td2F2 structure at a distance of 3.5 Å from the anomeric C1 atom of 2F-Glc, based on quantum mechanics/molecular mechanics metadynamics analysis (Fig. 5C in (30)). The side chain hydroxy group of Tyr295^B^ in Td2F2 is near the putative NW, and this conformation, along with Glu166, may support the nucleophilic attack by water during the deglycosylation step.

**Fig. 3.**
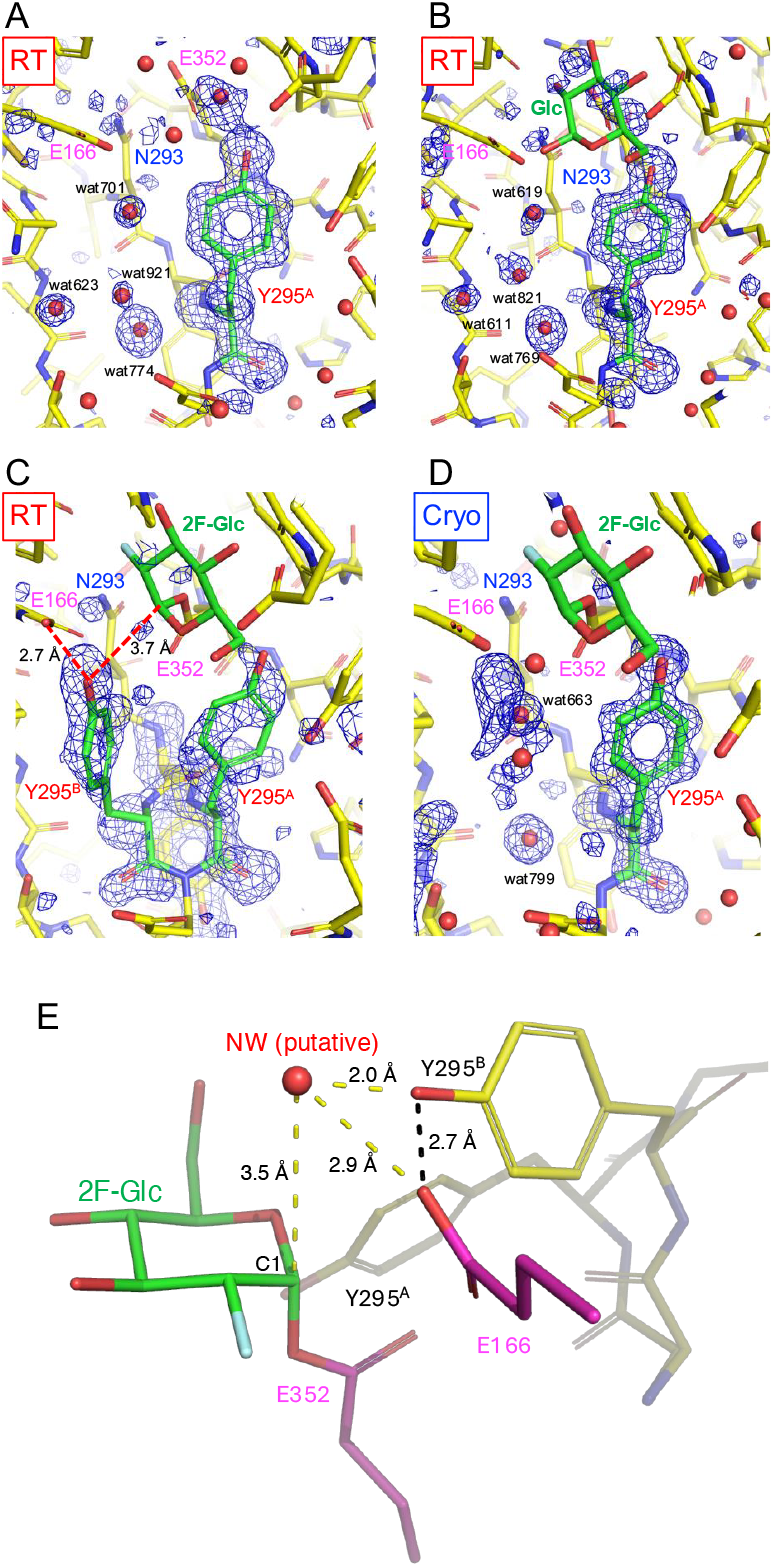
(*A*-*D*) Omit *mF*_o_-*DF*_c_ X-ray electron density (XRED) maps (blue mesh, 3.0σ) of Tyr295 and nearby water molecules, and (*E*) 2F-Glc complex structure with alternative conformations of Tyr295 and putative nucleophilic water (NW). (*A*) Ligand-free form at room temperature. (*B*) Glucose complex at room temperature. (*C*) 2F-Glc complex at room temperature with two alternative conformations, Tyr295^A^ and Tyr295^B^. (*D*) 2F-Glc complex at cryogenic temperature (100 K). Atoms of the following residues and water molecules were excluded from the map calculation: Tyr295, wat774, wat623, wat701, and wat921 in (*A*); Tyr295, wat769, wat611, wat619, and wat821 in (*B*); Tyr295^A^ and Tyr295^B^ in (*C*); and Tyr295, wat799, and wat663 atoms in (*D*). H and D atoms are not included in the models. (*E*) The putative NW is placed at a 3.5 Å distance from the anomeric C1 atom according to a quantum mechanics/molecular mechanics metadynamics analysis on the deglycosylation step of anomer-retaining GH129 3,6-anhydro-D-galactosidase (30). Atom-atom distances and hydrogen bonds are shown with yellow and black dotted lines, respectively.

### Kinetic analysis of the Y295F mutant and the catalytic role of tyrosine in the active center of GH1 and GH5

We constructed a Y295F mutant in which Tyr295 of Td2F2 was replaced by phenylalanine. Y295F had almost lost its activity toward disaccharides (*SI Appendix*, Table S8). The activity of Y295F toward the synthetic substrate pNP-Glc was also markedly reduced, and its kinetic parameters were measured (*SI Appendix*, Table S9). The *K*_m_ and *k*_cat_ values of the wild-type enzyme and Y295F mutant were 0.39 mM and 6.4 s^−1^, and 35 mM and 0.69 s^−1^, respectively. Therefore, the Y295F mutation resulted in an 820-fold reduction in *k*_cat_/*K*_m_, caused by a 90-fold increase in *K*_m_ and a 9.3-fold decrease in *k*_cat_. Interpreting these results according to the scheme shown below, where *E* corresponds to enzyme, *S* to substrate (pNP-Glc), *P*_1_ to aglycon *p*-nitrophenol, *ES\** to GEI, and *P*_2_ to glucose (31), the Y295F mutation resulted in a reduction in both *k*_1_ (Michaelis complex formation) and *k*_2_ (glycosylation step) rate constants. When glycosylation (GEI formation) is the rate-limiting step (*k*_2_ < *k*_3_), changes in *k*_3_ (deglycosylation step) do not significantly alter either *K*_m_ or *k*_cat_.

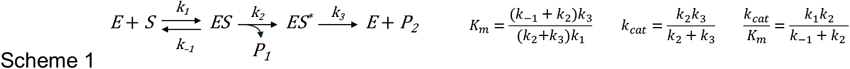

The catalytic role of the residue corresponding to Tyr295 in a GH1 enzyme has been demonstrated through detailed kinetic analyses of *Agrobacterium faecalis* BGL (*Af*BGL). Tyr298 in *Af*BGL was proposed to modulate the p*K*_a_ of the catalytic nucleophile Glu358 (32). In a catalytic nucleophile mutant of *Af*BGL (E358D), Tyr298 attacked the anomeric carbon of the substrate occasionally (1 time in 1,100) (33). Kinetic analysis of the Y298F mutant indicated that Tyr298 functions both in orienting the nearby nucleophile Glu358 and stabilizing the deprotonated state in the free enzyme. These conclusions, described in the 30-year-earlier studies using a GH1 BGL, align well with the structural features of Tyr295 in Td2F2 revealed in the present study. The role of tyrosine residue adjacent to the nucleophile residue has also been investigated in GH5 endoglucanase (RBcel1), whose active center is conserved with that of GH1. In RBcel1, GEI can be trapped using the Y201F mutant, suggesting that Tyr201 plays a critical role in the deglycosylation step (34).

## Conclusion

The proposed reaction mechanism of Td2F2, as elucidated in this study, is shown in Fig. 4. Throughout the reaction cycle, Glu166 is anchored by a hydrogen bond network involving Asn293, Asn165, and Arg77. H/D exchange data revealed that the acid/base catalyst Glu166 has a high proton transfer capacity. The catalytic nucleophile Glu352 undergoes a conformational change with the formation of a covalent intermediate, which in turn brings water molecules into the vicinity of the active center and moves the surrounding residues. In the catalysis of GH1 BGL, a tyrosine near the nucleophile plays key roles. Tyr295 in Td2F2 forms a hydrogen bond with Glu352 in the ligand-free structure (and presumably also in the Michaelis complex), supporting the Michaelis complex formation (*k*_1_) and glycosylation (*k*_2_) step, as suggested by the kinetic analysis. The alternative conformation of Tyr295 further suggests its involvement in the deglycosylation step (*k*_3_).

**Fig. 4.**
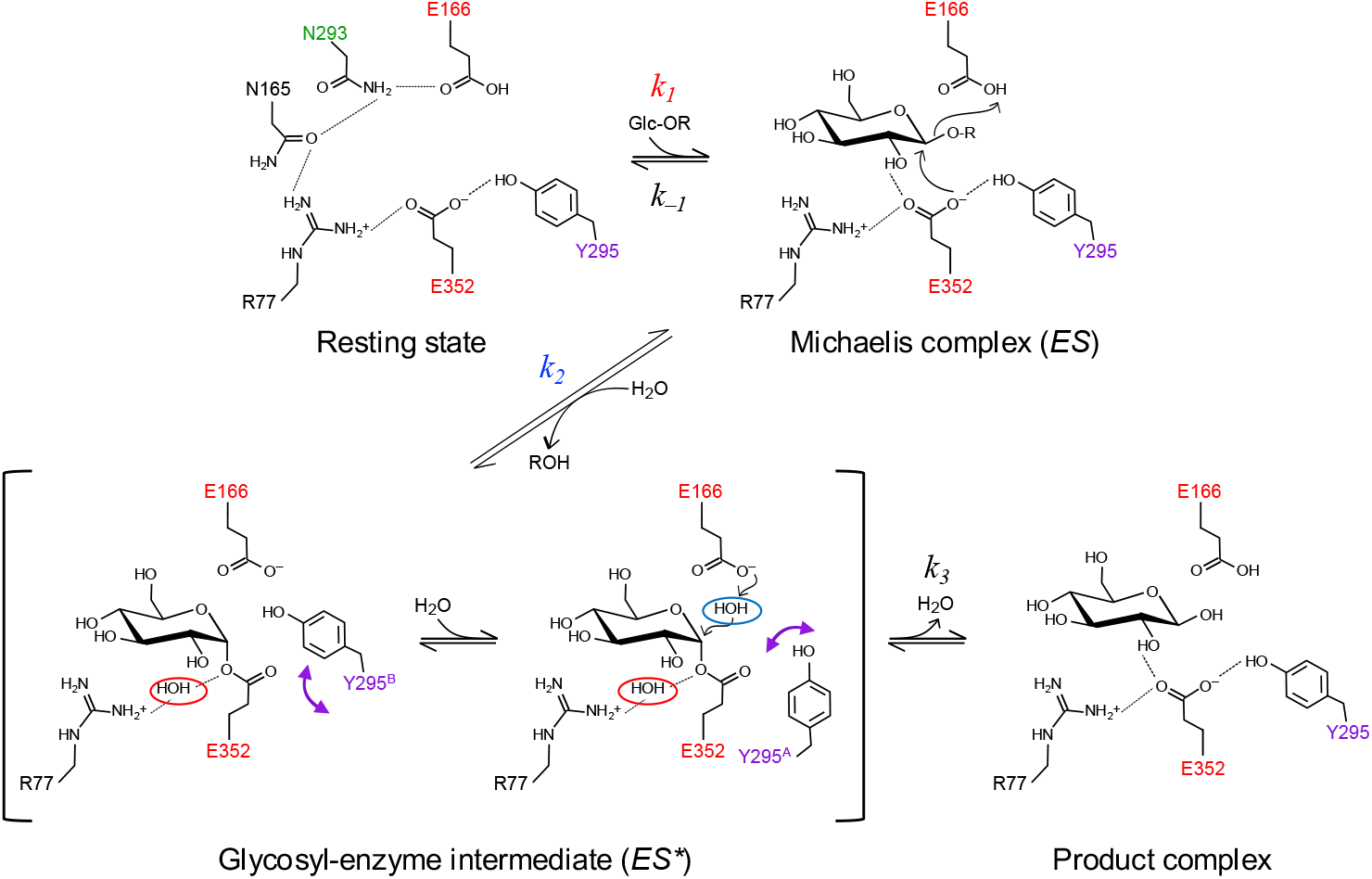
Proposed reaction mechanism of Td2F2. The acid/base catalyst Glu166 is fixed by a hydrogen bond network pivoted by Asn293. Tyr295 forms a hydrogen bond with the catalytic nucleophile Glu352 in the resting state, Michaelis complex, and product complex. In the glycosyl-enzyme intermediate, the side chain displacement of Glu352 induces water molecule incorporation (red circle) and conformational changes of Tyr295 (purple arrows). The nucleophilic water in the deglycosylation step is shown in a blue circle. Tyr295 plays an important role in the Michaelis complex formation (*k*_1_), glycosylation (*k*_2_), and deglycosylation (*k*_3_) steps.

In this study, we have determined the neutron structures for a representative anomer-retaining GH (GH1 BGL), including its covalent intermediate, at high resolution for the first time. In the covalent 2F-Glc complex, the hydrogen bonds involved in the hydroxy group recognition of the substrate were successfully determined in detail, including the positions of H/D atoms. A significant active site reorganization during the reaction cycle of GH1 BGL was observed. The importance of active site rearrangements in the glycosylation step has also been demonstrated in a computational analysis using a GH13 α-amylase (35). Crystallographic measurements at room temperature have the potential to observe important conformational changes that cannot be obtained by the widely used crystallographic or electron microscopic analyses under cryogenic conditions. Neutron structural analysis holds promise for future investigations into enzyme mechanisms.

## Materials and Methods

### Protein production and purification

The expression plasmid pJ-28a was constructed by replacing the multiple cloning site of pJexpress404 (DNA2.0) with that of pET-28a(+) (Novegen, Madison, WI, USA), incorporating the *td2f2* gene along with a C-terminal 6× histidine tag between NdeI and HindIII sites. The recombinant protein was expressed in *E. coli* Rosetta2 (DE3) (Sigma Aldrich Co., St. Louis, MO, USA). Transformants were cultured in 2× YT medium (Sigma-Aldrich) containing 100 μg/mL ampicillin at 37°C for 20 h. After centrifugation at 3,600 × *g* for 10 min, cells were cultured in lysogeny broth supplemented with 100 μg/mL ampicillin and 0.1 mM isopropyl-β-D-thiogalactopyranoside for 20 h at 37°C. Cells were harvested by centrifugation at 12,000 × *g* for 30 min, resuspended in 50 mM Tris-HCl buffer (pH 7.5), and lysed on ice using a sonicator. The lysates were incubated at 55°C for 30 min, followed by centrifugation at 32,300 × *g* for 40 min to obtain soluble proteins. The crude extract was purified using ammonium sulfate fractionation, immobilized metal affinity chromatography, and anion exchange chromatography. Ammonium sulfate was added to 30% saturation concentration (176 g/L), and the sample was centrifuged at 32,300 × *g* for 30 min to collect the supernatant. The supernatant was transferred to a dialysis tube with a molecular weight cutoff of 14,000 (Sekisui Material Solutions) and dialyzed against 50 mM Tris-HCl (pH 7.5). A Ni Sepharose 6 Fast Flow column (Cytiva, Malborough, MA, USA) was equilibrated with five times the volume of 50 mM Tris-HCl (pH 7.5) containing 5 mM imidazole and 0.3 M NaCl. After the addition of 5 mM imidazole and 0.3 M NaCl, the dialyzed sample was loaded onto the Ni column. The column was washed in 50 mM Tris-HCl (pH 7.5) containing 5 mM imidazole and 0.3 M NaCl, and Td2F2 was eluted using 50 mM Tris-HCl (pH 7.5) containing 300 mM imidazole and 0.3 M NaCl. The solution was collected and dialyzed in a dialysis tube with a molecular weight cutoff of 14,000 against 50 mM Tris-HCl (pH 7.5). Following dialysis, the sample was centrifuged at 3,700 × *g* for 10 min, and the supernatant was filtered using a Terumo syringe (Terumo, Tokyo, Japan) and a membrane filter (Advantec Toyo Kaisha Ltd., Tokyo, Japan). The protein sample was applied to a HiTrap Q XL column (Cytiva) equilibrated with 50 mM Tris-HCl (pH 7.5) and eluted using a linear gradient of 0 to 500 mM NaCl. The peak fraction, determined based on absorbance at 280 nm, was collected.

### Synthesis of DNP-2F-Glc

To prepare 2F-Glc complex protein, we synthesized high-purity DNP-2F-Glc (36) following previously reported procedures, via 1,3,4,6-tetra-*O*-acetyl-β-D-mannopyranose obtained from D-mannose as follows: 1,3,4,6-tetra-*O*-acetyl-β-D-mannopyranose was treated with dimethyaminosulfur trifluoride, yielding 3,4,6-tri-*O*-acetyl-2-fluoro-D-glucopyranose (37) electrospray ionization high-resolution mass spectroscopy (ESI HRMS) C_14_H_19_F_1_N_2_Na_1_O_9_ [M + Na]^+^ calcd for 373.0911, found 373.0894) followed by selective hydrolysis with (NH_4_)_2_CO_3_ in dimethylformamide afforded 3,4,6-tri-*O*-acetyl-2-fluoro-D-glucopyranose (ESI HRMS C_18_H_19_F_1_N_2_Na_1_O_12_ [M + Na]^+^ calcd for 497.0820, found 497.0806) in 83% yield over two steps. Next, glycoside formation reaction was performed on the resultant hemiacetal with 1-fluoro-2,4-dinitrobenzene in the presence of 1,4-diazabicyclo[2.2.2]octane and molecular sieves 4Å in DMF. Recrystallization from ethyl acetate and hexane yielded a desired 2,4-dinitrophenyl 3,4,6-tri-*O*-acetyl-2-fluoro-β-D-glucopyranoside (38) in 30% yield (α-isomer, 7%). The sample was then deacetylated under acidic conditions (MeOH−CHCl_3_−4N HCl in dioxane, 5:5:2) at room temperature, resulting in DNP-2F-Glc (ESI HRMS C_12_H_13_F_1_N_2_Na_1_O_12_ [M + Na]^+^ calcd for 371.0503, found 371.0502) with a 90% yield, which was used after recrystallization from methanol (39).

### Crystallization

Reagents dissolved in heavy water (99.9% D_2_O) were used for crystallization to facilitate H/D exchange of the protein and reduce incoherent background scattering from H atoms in the diffraction pattern. The purified protein solution was exchanged with 5 mM Tris pD 8.9 buffer using Amicon Ultracel-10K (Merck, Darmstadt, Germany) and concentrated to 20 or 30 mg/mL. Protein concentration was determined by measuring absorbance at 280 nm. A molar extinction coefficient of 100,520 M^−1^cm^−1^ was calculated using a theoretical extinction coefficient based on the amino acid sequence. A large-scale sitting-drop vapor diffusion method was adopted using a Falcon 60 × 15 mm center-well organ culture dish (Corning Inc., Corning, New York) (40). A siliconized glass cover slide was placed in the center well of the organ culture dish. A drop prepared by mixing 100 μL of protein solution and 100 μL of reservoir solution was placed on a cover slide. In the outside well of the culture dish, 2 mL of reservoir solution was poured. To reduce poly-crystallization of large crystals, 1-butylpyridinium chloride solution (No. 24) from the Ionic Liquid Screen Kit (Hampton Research, Aliso Viejo, CA, USA) was used. For ligand-free form crystals, a 20 mg/mL protein solution and reservoir solution consisting of 0.1 M CHES pD 9.5, 0.912 M K/Na tartrate•4H_2_O, 5% (w/v) 1-butylpyridinium chloride, and 0.2 M Li_2_SO_4_•H_2_O in heavy water were used. For glucose complex crystals, 20 mg/mL protein solution and reservoir solution consisting of 0.1 M CHES pD 9.5, 0.940 M K/Na tartrate•4H_2_O, 5% (w/v) 1-butylpyridinium chloride, 0.2 M Li_2_SO_4_•H_2_O, and 200 mM glucose in heavy water were used. Before crystallization, 30 mg/mL protein and 3 mM DNP-2F-Glc in 5 mM Tris pD 8.9 were incubated at 30°C for over 90 min. The color of the solution gradually changed from transparent to yellow. After the reaction, the protein solution was exchanged with 5 mM Tris pD 8.9 buffer using Amicon Ultracel-10K and concentrated to 30 mg/mL. For crystallization, this protein solution and reservoir solution consisting of 0.1 M CHES pD 9.5, 0.890 M K/Na tartrate•4H_2_O, 5% (w/v) 1-butylpyridinium chloride, and 0.2 M Li_2_SO_4_•H_2_O in heavy water were used. The culture dish was sealed with a lid using high-vacuum grease (Dow Toray Co., Ltd., Tokyo, Japan) and incubated at 20°C until the crystals grew to >1 mm^3^. A rotary evaporator and heavy water were used to facilitate the H/D exchange of the K/Na tartrate•4H_2_O solution.

### Neutron and X-ray diffraction experiments at room temperature for the joint XN crystallography

The crystals were sealed in a quartz capillary with a 3.5 mm ϕ and 0.01 mm thickness (Hilgenberg GmbH, Malsfeld, Germany) with a custom-made stainless-steel magnet base. Time-of-flight neutron diffraction experiments were performed using the BL03 IBARAKI biological crystal diffractometer (iBIX) in the Materials and Life Science Experimental Facility (MLF) of the Japan Proton Accelerator Research Complex (J-PARC, Tokai, Japan) (41, 42) with 34 two-dimensional position-sensitive detectors equipped with a scintillator sheet and wavelength-shifting fiber at room temperature (43). The accelerator power of the proton beam for the spallation neutron source was 600 kW, 730 or 800 kW, and 730 or 816 kW for the ligand-free form, glucose complex, and 2F-Glc complex, respectively. A neutron diffraction dataset was collected using a circular beam of 5 mm diameter with a selected neutron wavelength range of 2.28–6.19 Å for the ligand-free form and 1.86–5.76 Å for the glucose complex and 2F-Glc complex. The capillary sample was placed on a three-axis goniometer and exposed to 590,000; 708,750 or 620,000; and 630,880 or 564,390 pulsed neutrons for the ligand-free form; glucose complex; and 2F-Glc complex for each crystal orientation, respectively. In total, 33, 33, and 34 goniometer settings for the ligand-free form, glucose complex, and 2F-Glc complex, respectively, were selected to collect the entire dataset required for structure refinement. Incoherent neutron scattering data were collected from a 4.8-mm vanadium sphere using the same neutron wavelength range as the protein crystals. This procedure was performed to correct the variance in the detection efficiency of pixels within one detector, the difference in neutron beam intensities by wavelength, and the difference in detection efficiency by wavelength. Data reduction was performed using STARGazer 3.4.3, 3.8.2, and 3.9.0 (44), which employs a profile-fitting method for peak integration (45). Data statistics were calculated using the unit cell constants determined via X-ray diffraction.

After neutron diffraction data measurements, the same crystals were used for the synchrotron X-ray diffraction experiments on BL-5A at the Photon Factory of the High Energy Accelerator Research Institute (KEK, Tsukuba, Ibaraki, Japan). The crystals were exposed to an X-ray beam of 1.0 Å wavelength with a beam size of 200 μm × 200 μm at 10% of the maximum intensity at room temperature. Diffraction images of 180 frames were obtained using the oscillation method with 1.0º steps and 1.0-s exposure per frame for the ligand-free and glucose complex, or 0.5-s exposure per frame for the 2F-Glc complex. The diffraction images were processed using XDS (46), and statistics for the data collection were calculated using AIMLESS (47).

### Joint XN refinement

Joint XN refinement was performed using PHENIX 1.19.2_4158 (48) and Coot (49). The X-ray and neutron diffraction datasets were merged into an MTZ format file using the reflection tool in PHENIX. Five percent of the reflections common to both datasets were randomly assigned as a test dataset for cross-validation. The initial structure model was solved via the molecular replacement method using X-ray data and the atomic coordinates of wild-type Td2F2 in its ligand-free form (PDB ID: 3WH5), from which the solvent water molecules and ligands were removed.

After several cycles of atomic coordinate and temperature factor refinement using X-ray intensity data, oxygen atoms of water molecules and carbon and oxygen atoms of CHES were placed in the model, and the X-ray refinement was repeated. For the glucose complex and 2F-Glc complex, glucose and 2F-Glc were also added to the model, respectively. After several cycles of joint XN refinement, NSLD peaks, attributable to H and D atoms, were observed in the *mF*_o_-*DF*_c_ map. H and D atoms were placed in the model using phenix.readyset. The exchangeable hydrogen site atoms were treated as disordered (multiple) models of H and D atoms, with initial occupancies set at 0.5:0.5. Then, exchangeable hydrogen site atoms other than those in the main chains were removed from the atomic coordinates. The coordinates and temperature factors for all atoms were refined using this model. The occupancies of only exchangeable hydrogen atom sites were also refined. H and D atoms at exchangeable sites within protein side chains, solvent water molecules, and ligands were manually added to the atomic coordinates and refined. These atoms were added when a peak was observed in the *mF*_o_-*DF*_c_ NSLD map or when non-hydrogen atoms bonded to exchangeable hydrogens were observed in the XRED map. The protein solution was exchanged with a heavy water solution after protein purification, and the heavy water solution was also used for crystallization; therefore, water was regarded as D_2_O, and D atoms with an occupancy of 1.0 were manually added. The procedures for modeling and refinements were repeated until all observed H and D atoms were included in the model. We confirmed whether adding H and D atoms reduced residual densities in the *mF*_o_-*DF*_c_ NSLD map. When negative peaks were confirmed in D atoms of water molecules, the initial occupancies were set at 0.5, and the atomic coordinates, temperature factors, and occupancies were further refined. All molecular figures were generated using PyMOL (Schrödinger LLC, New York, NY, USA).

### Cryogenic crystallography of wild-type Td2F2 complexed with 2F-Glc

The purified protein solution was exchanged with 5 mM Tris pD 8.9 buffer and concentrated to 20 mg/mL, following the same procedure used for large crystals. A sitting-drop vapor diffusion method was adopted using a Falcon 60 × 15 mm center well organ culture dish. In the outside well of the plate, 2 mL of reservoir solution consisting of 0.1 M CHES pD 9.5, 0.935 M K/Na tartrate•4H_2_O, 5% (w/v) 1-butylpyridinium chloride, and 0.2 M Li_2_SO_4_•H_2_O in heavy water was poured. A drop prepared by mixing 100 μL of the protein solution and 100 μL of the reservoir solution was placed on a cover slide. The culture dish was sealed with a lid using high-vacuum grease and incubated at 20°C. Crystals were soaked in a reservoir solution supplemented with glycerol in heavy water. To minimize crystal damage, the glycerol concentration was gradually increased to 30% (v/v) in 5% increments. The crystals were then scooped with a cryoloop (Hampton Research) and cryo-cooled by dipping into liquid nitrogen. Synchrotron X-ray diffraction experiment was performed on BL-5A at the Photon Factory. The crystal was exposed to an X-ray beam of 1.0 Å wavelength with a beam size of 50 μm (h) × 75 μm (v) at 100% of the maximum intensity at cryogenic temperature (100 K). Diffraction images of 360 frames were obtained using the oscillation method with 1.0° steps and 1.0-s exposure per frame. Data processing and crystallographic refinement were performed using the same program applied in joint XN crystallography at room temperature. The initial structure model was solved via molecular replacement method using X-ray data and the atomic coordinates of the room-temperature 2F-Glc complex structure, from which the H and D atoms, solvent water molecules, and ligands were removed. After several cycles of atomic coordinate and temperature factor refinement using the X-ray intensity data, water molecules, Na^+^, CHES, and 2F-Glc were placed in the model, and the X-ray refinement was repeated.

### Site-directed mutant analysis

The Y295F mutant was generated via site-directed mutagenesis using the KOD One PCR Master Mix (TOYOBO Co. Ltd., Osaka, Japan) with the following primers (mutated codon underlined): 5′-GGCTGGGCGTCAATTAC<u>TTT</u>GAGCGCATGCGAGCCGTCG-3′ (forward) and 5′-CGACGGCTCGCATGCGCTC<u>AAA</u>GTAATTGACGCCCAGCC-3′ (reverse). The recombinant enzyme, expressed in *E. coli*, was purified and assayed following the same procedure as for the wild-type Td2F2. For the measurement of disaccharide hydrolysis, the enzyme reaction mixture (20 μL) contained 20 μg/mL enzyme, 10 mM disaccharide, and 100 mM Na-acetate (pH 5.5). After incubation at 65°C for 0, 2, 4, 6, 8, and 10 min, a 10 μL aliquot was sampled and mixed with 300 μL of pre-mixed coloring reagent from the LabAssay Glucose kit (FUJIFILM Wako Pure Chemical Corp., Osaka, Japan). After incubation for 5 min at 37°C, absorbance was measured at 505 nm using a microplate reader (BioTek Synergy H1; Agilent Technologies, Inc., Santa Clara, CA, USA). The absorbance was calibrated using a standard curve of 0.25–10 mM glucose. For the measurement of the kinetic constants, the enzyme reaction mixture (20 μL) contained 12.5 μg/mL enzyme, various concentrations of pNP-Glc, and 100 mM Na-acetate (pH 5.5). After incubation at 65°C for various periods, the reaction was stopped by adding 50 μL 1M NaHCO_3_, then a 65 μL aliquot was sampled to measure the absorbance. For wild-type Td2F2, 0.1, 0.15, 0.2, 0.25, 0.3, 0.4, 0.5, 1, and 3 mM pNP-Glc was used, and the reaction was stopped at 0, 10, 20, 30, 40, 50, 60, and 70 s for 0.1–0.5 mM pNP-Glc, and at 0, 0.5, 1.0, 1.5, 2.0, 2.5, 3.0, 3.5 min for 1 and 3 mM pNP-Glc. For Y295F mutant, 10, 15, 30, 35, 50, 70, and 100 mM pNP-Glc was used, and the reaction was stopped at 0, 20, 40, 60, 80, 100, 120, and 140 min for 10–35 mM pNP-Glc, and at 0, 10, 20, 30, 40, 50, 60, and 70 min for 50–100 mM pNP-Glc. Absorbance was measured at 405 nm using a microplate reader (BioTek Synergy H1) and calibrated using a standard curve of 0.05–1 mM *p*-nitrophenol. For all enzyme assays, triplicates were measured at each time point.

## Acknowledgments

We would like to thank the staff of the Structural Biology Research Center and Photon Factory at KEK for X-ray data collection, the staff of iBIX and MLF at J-PARC for neutron data collection, Dr. Michiko Konno and Dr. Taro Yamada for their support with neutron crystallography, and Dr. Katsuro Yaoi, Dr. Taku Uchiyama, and Dr. Tomohiko Matsuzawa for providing the *td2f2* gene, and Dr. Takatoshi Arakawa, Dr. Chihaya Yamada, Dr. Akimasa Miyanaga, and Dr. Toma Kashima for helpful discussions. The neutron experiment was conducted as a project for the Ibaraki Prefectural Local Government Beam Line at J-PARC MLF (Proposal Nos. 2020PX2011 and 2022PX3004). The X-ray experiments were conducted with the approval of the Photon Factory Program Advisory Committee (Proposal Nos. 2020G021, 2021G052, 2022G034, and 2023G052). This study was partially supported by JSPS-KAKENHI (22K06157 to N.Y., and 19H00929 and 24H02269 to S.F. and A.I.).

