## Supporting Information for "Neutron crystallography of the covalent intermediate of β-glucosidase reveals remodeling of the catalytic center"

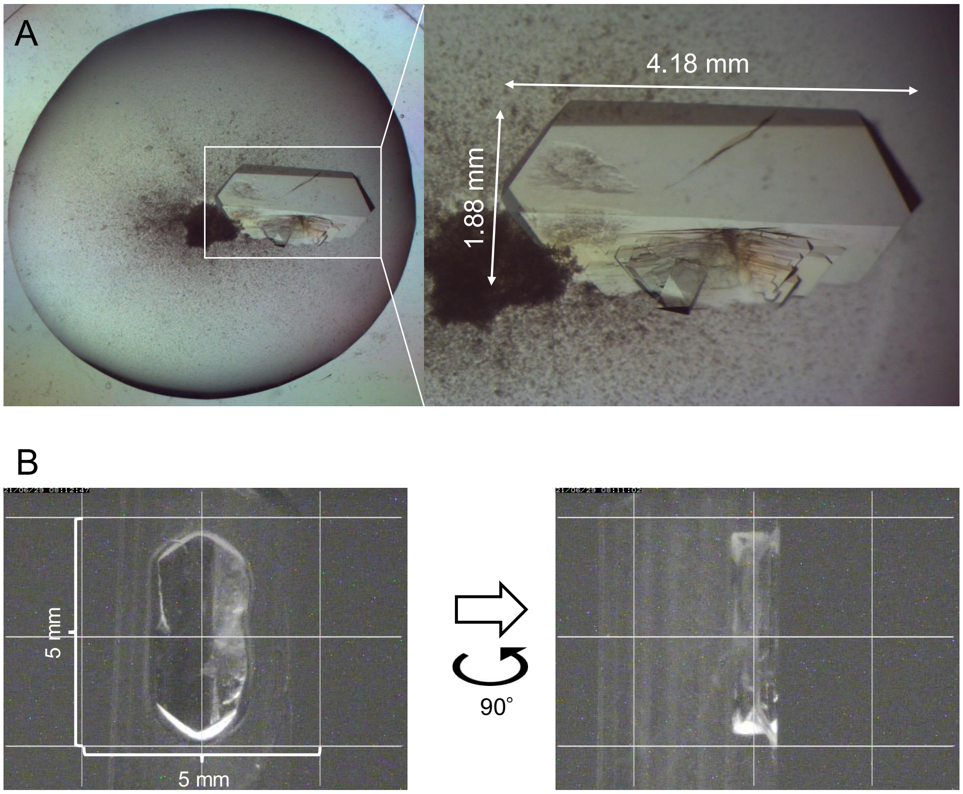


**Fig. S1.** Photographs of the large crystal in ligand-free form. (*A*) The crystallization drop. The initial drop volume was 200 μL. The crystal used for the diffraction experiments is squared. (*B*) The crystal was sealed in a quartz capillary.

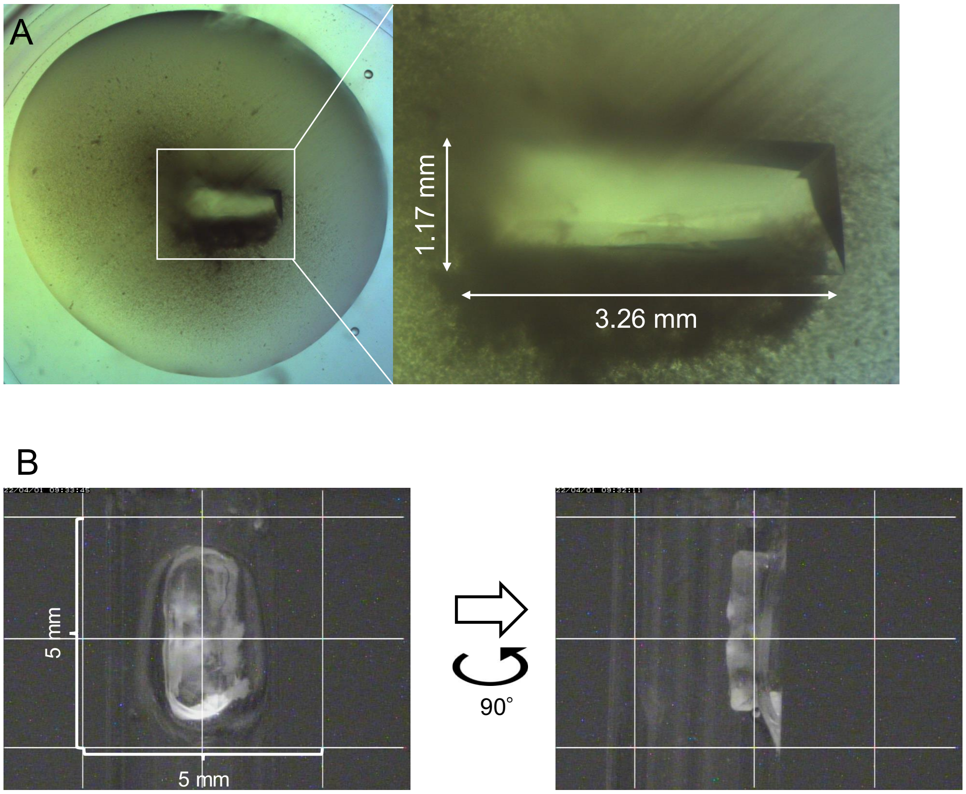


**Fig. S2.** Photographs of the large crystal of the glucose complex. (*A*) The crystallization drop. The initial drop volume was 200 μL. The crystal used for the diffraction experiments is squared. (B) The crystal was sealed in a quartz capillary.

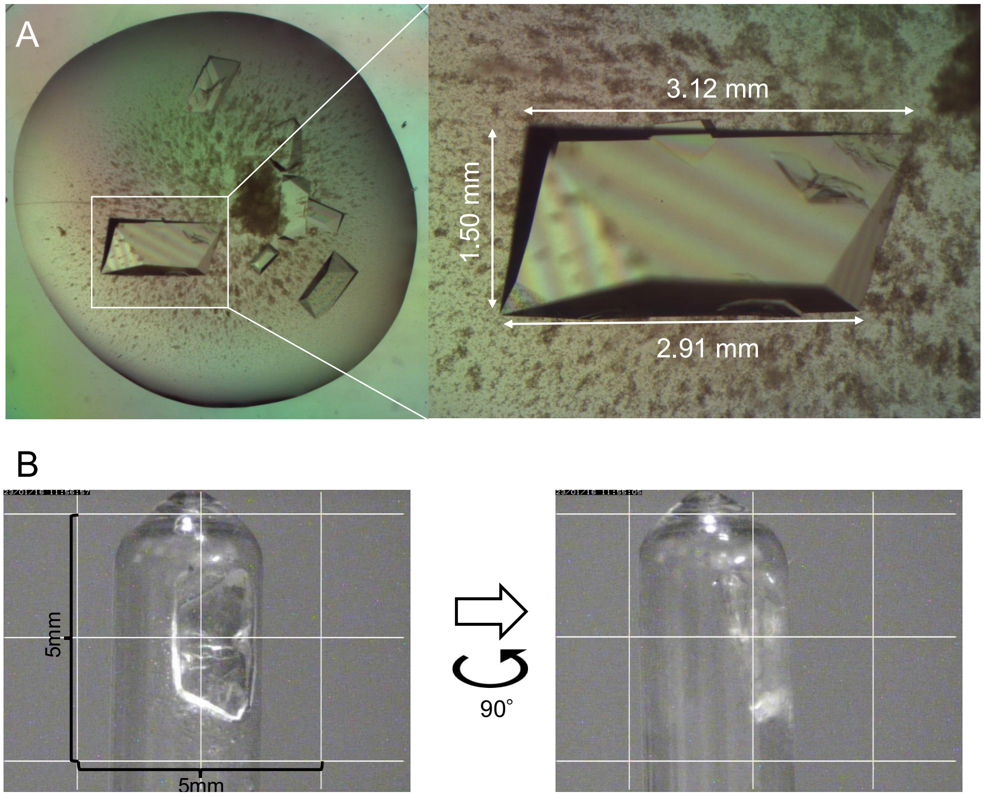


**Fig. S3.** Photographs of the large crystal of the 2F-Glc complex. (*A*) The crystallization drop. The initial drop volume was 200 μL. The crystal used for the diffraction experiments is squared. (*B*) The crystal was sealed in a quartz capillary.

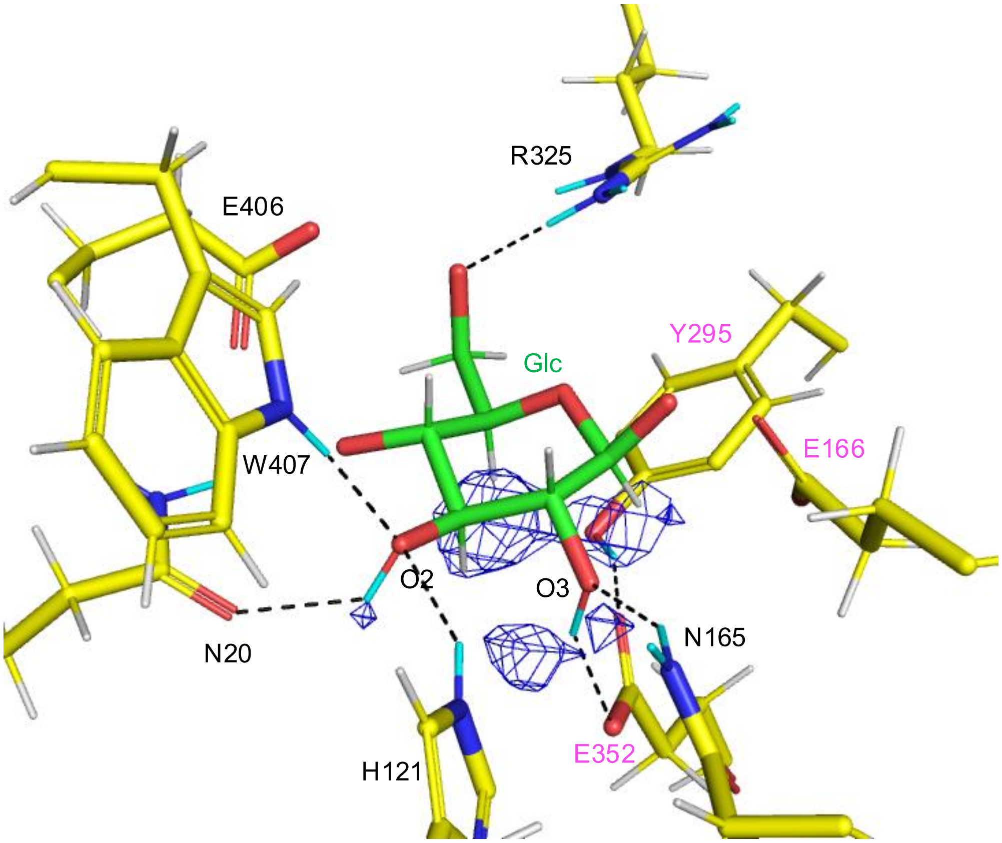


**Fig. S4.** An $mF_o$-$DF_c$ neutron scattering length density (NSLD) map (blue mesh, 2.7σ) in the glucose complex structure. H and D atoms are colored white and cyan, respectively. The following atoms were excluded from the map calculation: DO2 and DO3 of Glc.

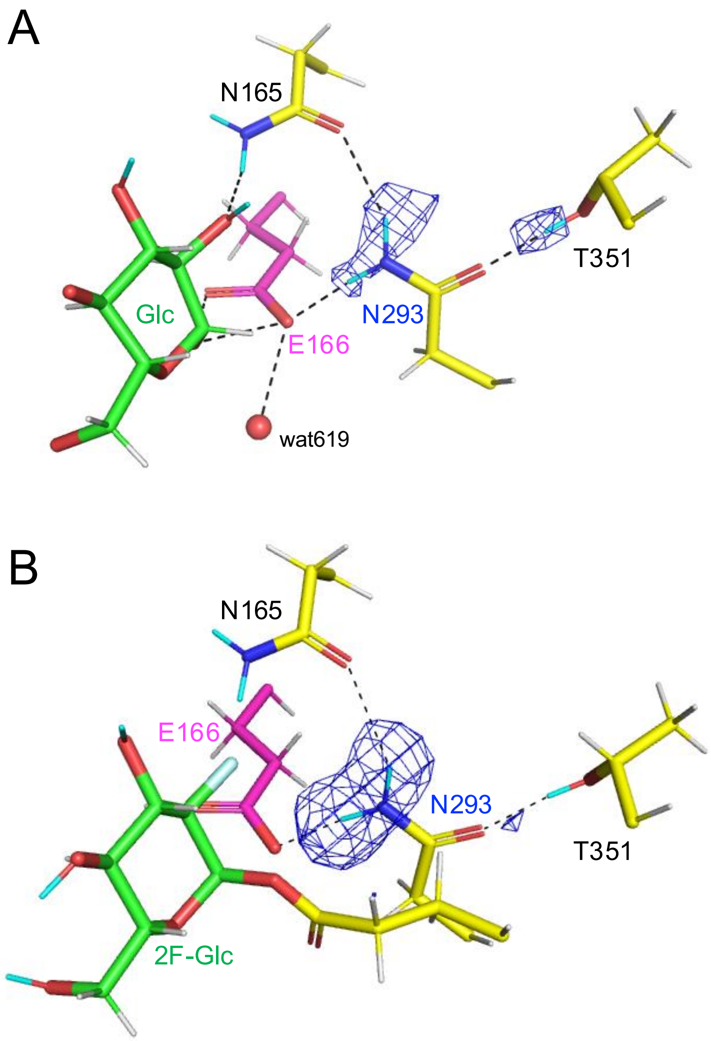


**Fig. S5.** Asn293 and surrounding residues with $mF_o$-$DF_c$ NSLD maps (blue mesh). (*A*) Glucose complex structure (3.3σ). (*B*) 2F-Glc complex structure (2.7σ). H and D atoms are colored white and cyan, respectively. The following atoms were excluded from the map calculation: DD21 and DD22 of Asn293 and DG1 of Thr351.

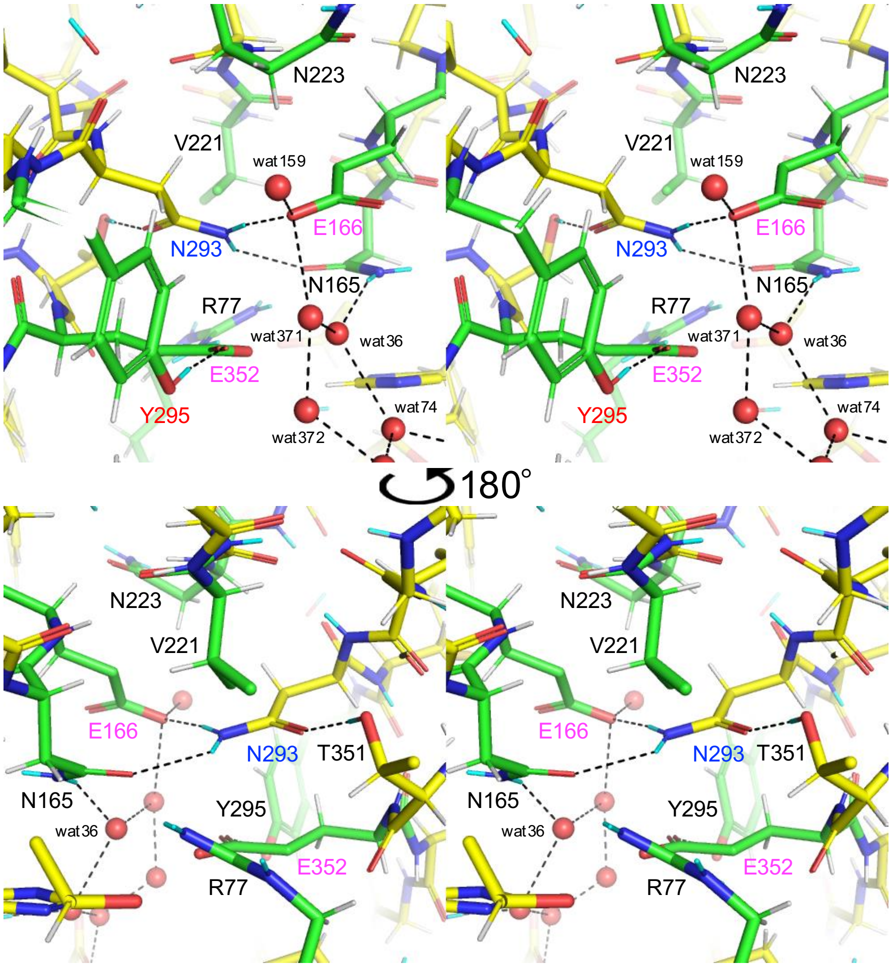


**Fig. S6.** Stereo views of Asn293 and surrounding residues in the ligand-free structure. H and D atoms are colored white and cyan, respectively. Hydrogen bonds are shown with black dotted lines. Views from both sides are shown (top and bottom).

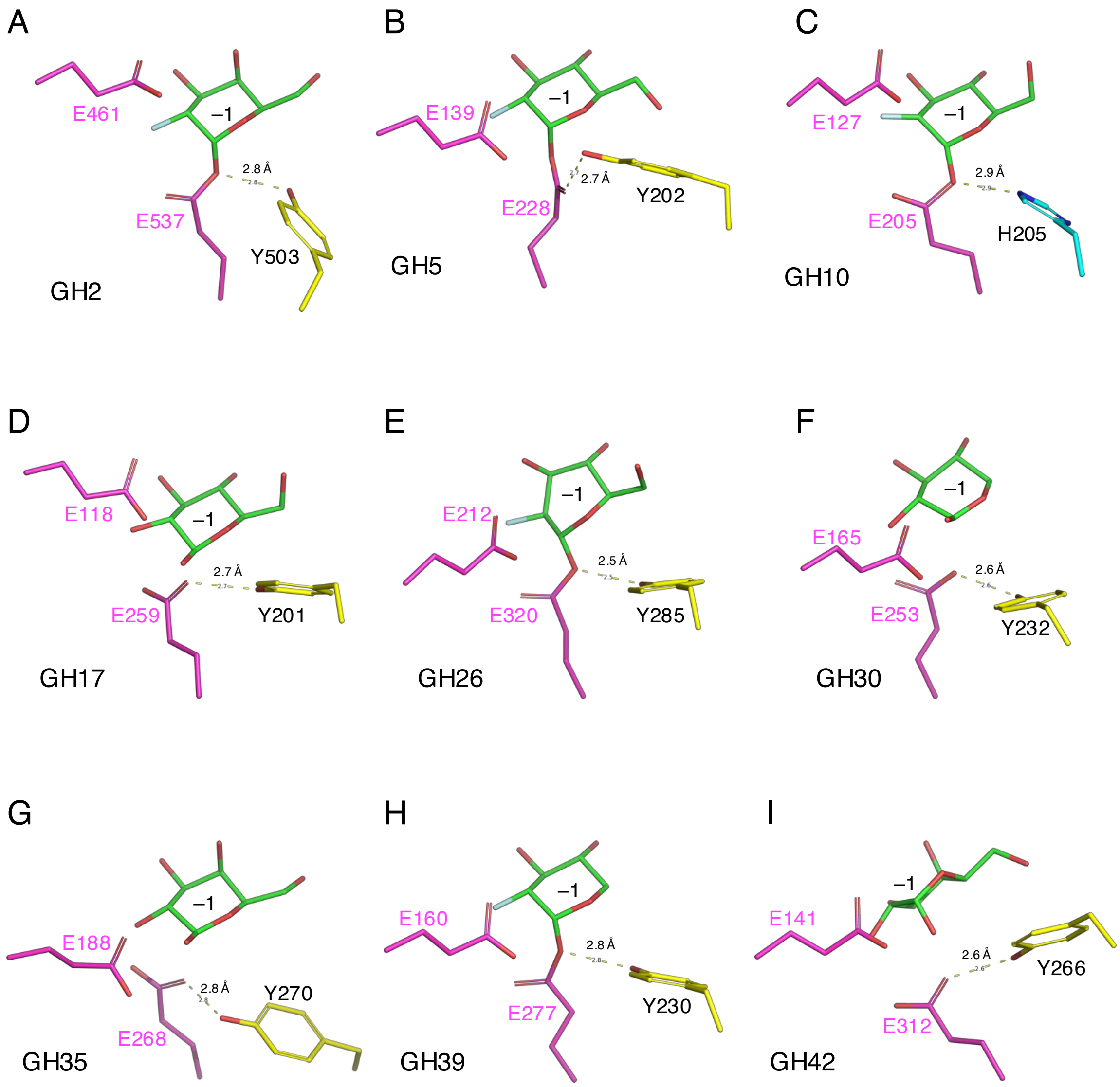


**Fig. S7.** Active center structure of clan GH-A enzymes. (*A*) *Escherichia coli* GH2 β-galactosidase LacZ (PDB ID: 1JZ2) (1). (*B*) *Bacillus agaradhaerens* GH5 endoglucanase Cel5A (PDB ID: 1H11) (2). (*C*) *Cellulomonas fimi* GH10 xylanase B (PDB ID: 1EXP) (3). (*D*) Potato GH17 endo-1,3-β-glucanase (PDB IDs: 3UR8 and 4GZJ) (4, 5). (*E*) *Pseudomonas cellulosa* GH26 β-mannanase (PDB IDs: 1GW1 and 1J9Y) (6, 7). (*F*) *Erwinia chrysanthemi* GH30 glucuronoxylanase (PDB ID: 2Y24) (8). (*G*) Human GH35 β-galactosidase (PDB ID: 3THC) (9). (*H*) *Thermoanaerobacterium saccharolyticum* GH39 β-xylosidase (PDB ID: 1UHV) (10). (*I*) *Thermus thermophilus* A4 GH42 β-galactosidase (PDB ID: 1KWK) (11). The catalytic nucleophile and acid/base catalyst residues (magenta), nearby tyrosine (yellow) or histidine (blue), and a sugar moiety at subsite –1 (green) are shown.

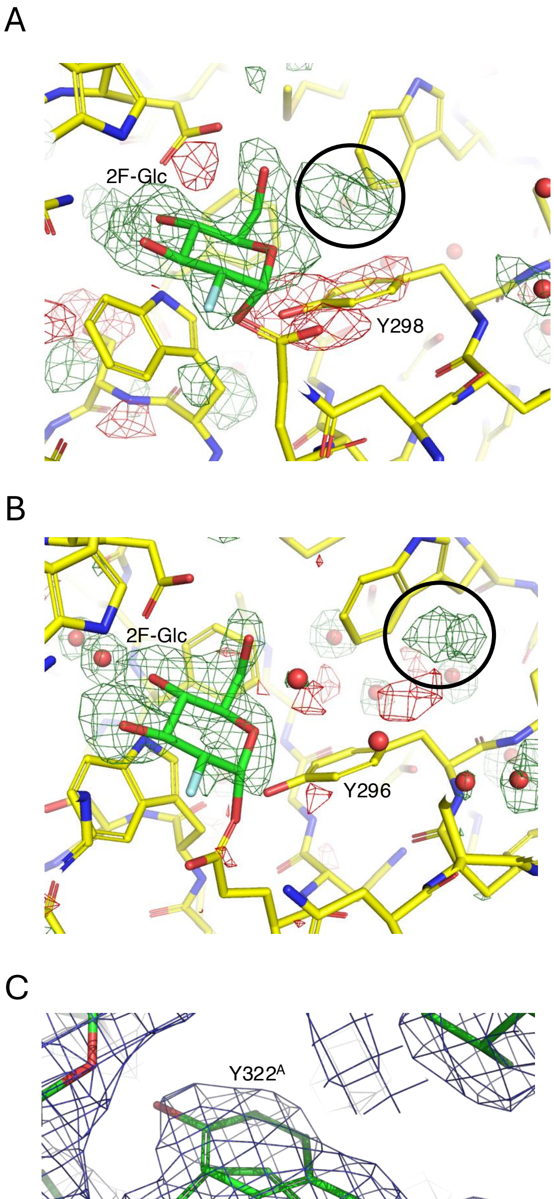


**Fig. S8.** Electron density maps of Tyr295 equivalent residues in X-ray crystallographic 2F-Glc complex structures of GH1 BGLs. (*A*) *P. polymyxa* BGL with $mF_o$-$DF_c$ X-ray electron density (XRED) map (green for +3.0σ and red for –3.0σ) (PDB ID: 2JIE, 2.3 Å resolution at 120 K). (*B*) *H. orenii* BGL with $mF_o$-$DF_c$ XRED map (green for +3.0σ and red for –3.0σ) (PDB ID: 4PTW, 2.0 Å resolution at 100 K). (*C*) *O. sativa* BGL Os4BGlu12 with $2mF_o$-$DF_c$ XRED map (1.0σ, blue mesh) (PDB ID: 3PTM, 2.4 Å at 100 K). In (*A*) and (*B*), 2F-Glc was excluded from the map calculation, and a density peak at a position corresponding to $Tyr295^B$ in Td2F2 is circled (black).

**Table S1.** Crystallographic data collection and refinement statistics for the ligand-free form at room temperature.

| Dataset | Td2F2 ligand-free form (neutron, room temperature) | |
|---|---|---|
| **Data collection** | | |
| | X-ray | Neutron |
| Beamline | PF BL-5A | J-PARC MLF BL03 |
| Temperature | Room temperature | Room temperature |
| Wavelength (Å) | 1.0000 | 2.28–6.19 |
| Space group | $P2_12_12_1$ | |
| Unit cell (Å) | *a = 69.72, b = 71.03, c = 97.29* | |
| Resolution (Å) | 44.30–1.19 (1.21–1.19) | 22.15–1.80 (1.86–1.80) |
| Total reflections | 1,011,695 (48,114) | 362,071 (27,623) |
| Unique reflections | 154,133 (7,472) | 45,104 (4,400) |
| Completeness (%) | 99.6 (98.6) | 99.1 (98.3) |
| Redundancy | 6.6 (6.4) | 8.0 (6.3) |
| Mean $I/\sigma(I)$ | 11.4 (2.1) | 8.6 (1.7) |
| $R_{merge}$ (%) | 7.9 (82.9) | 25.0 (97.2) |
| $R_{pim}$ (%) | 3.3 (34.7) | 9.3 (41.0) |
| $CC_{1/2}$ | 0.998 (0.755) | 0.983 (0.466) |
| Wilson B-factor (Å$^2$) | 11.44 | |
| **Refinement** | | |
| Resolution (Å) | 44.30–1.19 (1.20–1.19) | 22.15–1.80 (1.84–1.80) |
| No. of reflections | 154,041 | 45,096 |
| $R_{work}$ (%) | 15.2 (25.2) | 14.9 (27.6) |
| $R_{free}$ (%) | 16.8 (23.9) | 17.3 (31.3) |
| Number of atoms | 7,723 | |
| RMSD from ideal values | | |
| Bond lengths (Å) | 0.009 | |
| Bond angles (°) | 1.003 | |
| Ramachandran plot (%) | | |

| | |
|---|---|
| Favored | 97.5 |
| Allowed | 2.5 |
| Outlier | 0 |
| PDB code | 9LPH |

Unit cell parameters were determined using X-ray diffraction data. Values in parentheses represent the highest resolution shell.

**Table S2.** Crystallographic data collection and refinement statistics for the glucose complex form at room temperature.

| Dataset | Td2F2 + glucose (neutron, room temperature) | |
|---|---|---|
| **Data collection** | | |
| | X-ray | Neutron |
| Beamline | PF BL-5A | J-PARC MLF BL03 |
| Temperature | Room temperature | Room temperature |
| Wavelength (Å) | 1.0000 | 1.86–5.76 |
| Space group | $P2_12_12_1$ | |
| Unit cell (Å) | *a = 69.65, b = 71.09, c = 97.38* | |
| Resolution (Å) | 44.30–1.20 (1.22–1.20) | 19.74–1.70 (1.76–1.70) |
| Total reflections | 971,505 (48,046) | 380,254 (26,556) |
| Unique reflections | 144,269 (6,851) | 52,857 (5,058) |
| Completeness (%) | 95.7 (92.7) | 98.1 (95.7) |
| Redundancy | 6.7 (7.0) | 7.2 (5.3) |
| Mean $I/\sigma(I)$ | 14.2 (2.2) | 7.9 (1.7) |
| $R_{merge}$ (%) | 7.2 (88.5) | 22.4 (82.8) |
| $R_{pim}$ (%) | 3.0 (35.5) | 8.8 (37.8) |
| $CC_{1/2}$ | 0.998 (0.730) | 0.984 (0.428) |
| Wilson B-factor (Å$^2$) | 11.01 | |
| **Refinement** | | |
| Resolution (Å) | 44.30–1.20 (1.21–1.20) | 19.74–1.70 (1.73–1.70) |
| No. of reflections | 144,184 | 52,854 |
| $R_{\mathrm{work}}$ (%) | 15.3 (25.3) | 14.8 (26.5) |
| $R_{\mathrm{free}}$ (%) | 16.2 (28.3) | 17.0 (29.5) |
| Number of atoms | 7,851 | |
| RMSD from ideal values | | |
| Bond lengths (Å) | 0.007 | |
| Bond angles (°) | 1.210 | |

| | |
|---|---|
| Ramachandran plot (%) | |
| Favored | 97.3 |
| Allowed | 2.7 |
| Outlier | 0 |
| PDB code | 9LPI |

Unit cell parameters were determined using X-ray diffraction data. Values in parentheses represent the highest resolution shell.

**Table S3.** Crystallographic data collection and refinement statistics for the 2F-Glc complex form at room temperature.

| Dataset | Td2F2 + 2F-Glc (neutron, room temperature) | |
|---|---|---|
| **Data collection** | | |
| | X-ray | Neutron |
| Beamline | PF BL-5A | J-PARC MLF BL03 |
| Temperature | Room temperature | Room temperature |
| Wavelength (Å) | 1.0000 | 1.86–5.76 |
| Space group | $P2_12_12_1$ | |
| Unit cell (Å) | *a = 69.54, b = 71.23, c = 97.20* | |
| Resolution (Å) | 44.29–1.32 (1.34–1.32) | 20.07–1.70 (1.76–1.70) |
| Total reflections | 736,840 (35,174) | 400,573 (28,259) |
| Unique reflections | 110,952 (5,330) | 53,007 (5,139) |
| Completeness (%) | 97.7 (95.6) | 98.5 (97.0) |
| Redundancy | 6.6 (6.6) | 7.6 (5.5) |
| Mean $I/\sigma$ ($I$) | 13.5 (2.1) | 10.6 (2.0) |
| $R_{merge}$ (%) | 7.8 (84.3) | 18.4 (76.0) |
| $R_{pim}$ (%) | 3.2 (35.1) | 7.1 (34.3) |
| $CC_{1/2}$ | 0.998 (0.746) | 0.991 (0.556) |
| Wilson B-factor (Å$^2$) | 14.00 | |
| **Refinement** | | |
| Resolution (Å) | 43.29–1.32 (1.33–1.32) | 20.07–1.70 (1.73–1.70) |
| No. of reflections | 110,877 | 53,005 |
| $R_{work}$ (%) | 14.9 (26.9) | 15.5 (25.1) |
| $R_{free}$ (%) | 16.0 (29.0) | 17.0 (28.7) |
| Number of atoms | 7,920 | |
| RMSD from ideal values | | |
| Bond lengths (Å) | 0.010 | |
| Bond angles (°) | 1.054 | |

| | |
|---|---|
| Ramachandran plot (%) | |
| Favored | 97.3 |
| Allowed | 2.7 |
| Outlier | 0 |
| PDB code | 9LPV |

Unit cell parameters were determined using X-ray diffraction data. Values in parentheses represent the highest resolution shell.

**Table S4.** Occupancy and temperature factors of D:H and water atoms in the active site.

| | Ligand-free | Glucose | 2F-Glc |
|---|---|---|---|
| Asn293 | | | |
| DD21:HD21 | 0.71 (17.49):0.29 (17.49) | 0.60 (17.08):0.39 (17.08) | 0.85 (13.58):0.15 (13.58) |
| DD22:HD22 | 0.75 (17.49):0.25 (17.49) | 0.60 (17.30):0.40 (17.30) | 0.89 (13.58):0.11 (13.58) |
| Tyr295 | | | |
| DH:HH | 0.53 (18.87):0.47 (18.87) | 0.66 (16.61):0.34 (16.85) | – |
| Glc/2F-Glc[a] | | | |
| DO2:HO2 | – | 0.54 (19.81):0.46 (19.79) | – |
| DO3:HO3 | – | 0.52 (18.84):0.47 (19.19) | 0.71 (12.26):0.29 (12.26) |
| DO4:HO4 | – | – | 0.63 (12.86):0.37 (12.86) |
| DO6:HO6 | – | – | 0.74 (17.24):0.26 (17.24) |
| Waters | | | |
| Wat648 O, D1, D2 | – | 1.00 (17.46), 0.38 (17.93), 0.36 (16.57) | 1.00 (10.45), 0.64(12.54), 0.64(12.54) |
| Wat653 O | 1.00 (18.86) | – | – |
| Wat734 O, D1, D2 | – | – | 1.00 (17.43), 0.42 (20.91), 0.39 (20.91) |
| Wat644 O | 1.00 (23.53) | – | – |
| Wat701 O | 1.00 (32.66) | – | – |
| Wat853 O | 1.00 (31.39) | – | – |
| Wat824 O | 1.00 (38.45) | – | – |
| Wat621 O | 1.00 (27.14) | – | – |

Values in parentheses are the temperature factors ($Å^2$). The occupancy of water atoms was set to 1.0. [a] Atoms in glucose or 2F-Glc.

**Table S5.** Refinement statistics for the 2F-Glc complex form (X-ray structure) at room temperature.

| Dataset | Td2F2 + 2F-Glc (X-ray, room temperature) |
|---|---|
| **Refinement** | |
| Resolution (Å) | 44.29–1.32 (1.33–1.32) |
| No. of reflections | 110,877 |
| $R_{work}$ (%) | 15.7 (26.0) |
| $R_{free}$ (%) | 16.8 (28.7) |
| Number of atoms | 3,859 |
| Number of waters | 308 |
| RMSD from ideal values | |
| Bond lengths (Å) | 0.005 |
| Bond angles (°) | 0.897 |
| Average B-factor (Å$^2$) | |
| Overall | 19.75 |
| Protein | 18.82 |
| Water | 30.62 |
| Ramachandran plot (%) | |
| Favored | 97.3 |
| Allowed | 2.7 |
| Outlier | 0 |
| PDB code | 9LPX |

The X-ray diffraction dataset shown in Table S3 was used for refinement. Values in parentheses represent the highest resolution shell.

**Table S6.** Side-chain conformations of Tyr295 equivalent residues in X-ray crystallographic 2F-Glc complex structures of GH1 BGLs and myrosinase.

| PDB ID | Temp. (K)[a] | Resol. (Å)[b] | Tyr no.[c] | Enzyme | Note |
|---|---|---|---|---|---|
| 1E4I | 100 | 2.00 | 296 | *Paenibacillus polymyxa* BGL | No structure factor. Equal to 2MYR. |
| 1E70 | 100 | 1.65 | 330 | *Sinapis alba* myrosinase | No multi-conformation density. |
| 1E73 | 100 | 1.50 | 330 | *Sinapis alba* myrosinase | No multi-conformation density. Blocked by ascorbate. |
| 1OIN | 100 | 2.15 | 295 | *Thermotoga maritima* BGL | No multi-conformation density. |
| 1UWS | 100 | 1.95 | 322 | *Sulfolobus solfataricus* BGL | No multi-conformation density. |
| 1UYQ | 120 | 2.10 | 296 | *Paenibacillus polymyxa* BGL | No multi-conformation density. |
| 2RGM | 100 | 1.55 | 315 | *Oryza sativa* BGL Os3BGlu7 | No multi-conformation density. Blocked by glycerol. |
| 2JIE | 120 | 2.30 | 298 | *Paenibacillus polymyxa* BGL | Slight electron density peak at Y295 (B). See Fig. S8*A.* |
| 3AHV | 105 | 1.89 | 315 | *Oryza sativa* BGL Os3BGlu7 | No multi-conformation density. Blocked by glycerol. Equal to 3F5I. |
| 3AIR | 95 | 2.00 | 334 | *Triticum aestivum* BGL | No multi-conformation density. Blocked by 2,4-dinitrophenol group (aglycone). |
| 3AIW | 95 | 2.40 | 334 | *Secale cereale* BGL | No multi-conformation density. Blocked by 2,4-dinitrophenol group (aglycone). |
| 3GNR | 110 | 1.81 | 321 | *Oryza sativa* BGL Os3BGlu6 | No multi-conformation density. |
| 3PTM | 100 | 2.40 | 322 | *Oryza sativa* BGL Os4BGlu12 | Multiple conformations with slight rotation (30°) of $\chi_1$ (Cα-Cβ) angle. The density for an alternative conformation is ambiguous. Blocked by glycerol. See Fig. S8*C.* |
| 4PTW | 100 | 2.00 | 298 | *Halothermothrix orenii* BGL | Slight electron density peak at Y295 (B). See Fig. S8*B.* |

[a] Crystallographic data collection temperature. [b] Crystallographic resolution. [c] Residue number of Tyr corresponding to Y295 in Td2F2.

**Table S7.** Crystallographic data collection and refinement statistics for the 2F-Glc complex form (X-ray structure) at cryogenic temperature.

| Dataset | Td2F2 + 2F-Glc (X-ray, cryogenic temperature) |
|---|---|
| **Data collection** | X-ray |
| Beamline | PF BL-5A |
| Temperature (K) | 100 |
| Wavelength (Å) | 1.0000 |
| Space group | $P2_12_12_1$ |
| Unit cell (Å) | *a = 68.73, b = 69.79, c = 95.99* |
| Resolution (Å) | 48.97–1.11 (1.13–1.11) |
| Total reflections | 2,331,801 (104,499) |
| Unique reflections | 182,060 (8,940) |
| Completeness (%) | 100.0 (100.0) |
| Redundancy | 12.8 (11.7) |
| Mean $I/\sigma$ ($I$) | 12.9 (2.1) |
| $R_{merge}$ (%) | 16.0 (129.5) |
| $R_{pim}$ (%) | 4.6 (39.2) |
| $CC_{1/2}$ | 0.999 (0.724) |
| Wilson B-factor (Å$^2$) | 7.19 |
| **Refinement** | |
| Resolution (Å) | 48.97–1.11 (1.12–1.11) |
| No. of reflections | 181,940 |
| $R_{work}$ (%) | 16.4 (23.8) |
| $R_{free}$ (%) | 17.6 (23.5) |
| Number of atoms | 4,020 |
| Number of waters | 400 |
| RMSD from ideal values | |
| Bond lengths (Å) | 0.005 |
| Bond angles (°) | 0.883 |
| Average B-factor (Å$^2$) | |
| Overall | 12.95 |
| Protein | 11.70 |
| Water | 24.02 |
| Ramachandran plot (%) | |
| Favored | 97.3 |
| Allowed | 2.7 |
| Outlier | 0 |
| PDB code | 9LPY |

Values in parentheses represent the highest resolution shell.

**Table S8.** Hydrolytic activity (μmol $min^{-1}$ $mg^{-1}$ enzyme) of the wild-type and Y295F mutant of Td2F2 toward disaccharides.

| Substrate | Wild-type | Y295F |
|---|---|---|
| Sophorose (Glc-β1,2-Glc) | 28.5 ± 1.0 | ND |
| Laminaribiose (Glc-β1,3-Glc) | 20.6 ± 0.4 | ND |
| Cellobiose (Glc-β1,4-Glc) | 17.1 ± 0.8 | ND |
| Gentiobiose (Glc-β1,6-Glc) | 5.3 ± 0.1 | ND |

Activity was measured in 100 mM Na-acetate (pH 5.5) with 10 mM substrate at 65°C. ND, not detected (<0.4 μmol $min^{-1}$ $mg^{-1}$ enzyme).

**Table S9.** Kinetic parameters for pNP-Glc with Td2F2 and the Y295F mutant.

| | $K_m$ (mM) | $k_{cat}$ ($s^{-1}$) | $k_{cat}/K_m$ ($s^{-1}$ $mM^{-1}$) |
|---|---|---|---|
| Wild type | 0.39 ± 0.04 | 6.4 ± 0.3 | 16.4 ± 1.8 |
| Y295F | 35 ± 3 | 0.69 ± 0.03 | 0.02 ± 0.002 |
